# Simulating Pedigrees Ascertained for Multiple Disease-Affected Relatives

**DOI:** 10.1101/234153

**Authors:** Christina Nieuwoudt, Samantha J. Jones, Angela Brooks-Wilson, Jinko Graham

## Abstract

**Background:** Studies that ascertain families containing multiple relatives affected by disease can be useful for identification of causal, rare variants from next-generation sequencing data.

**Results:** We present the R package `SimRVPedigree`, which allows researchers to simulate pedigrees ascertained on the basis of multiple, affected relatives. By incorporating the ascertainment process in the simulation, `SimRVPedigree` allows researchers to better understand the within-family patterns of relationship amongst affected individuals and ages of disease onset.

**Conclusions:** Through simulation, we show that affected members of a family segregating a rare disease variant tend to be more numerous and cluster in relationships more closely than those for sporadic disease. We also show that the family ascertainment process can lead to apparent anticipation in the age of onset. Finally, we use simulation to gain insight into the limit on the proportion of ascertained families segregating a causal variant. `SimRVPedigree` should be useful to investigators seeking insight into the family-based study design through simulation.

## Background

Family-based studies of pedigrees with multiple disease-affected relatives are regaining traction for identification of rare causal variants. These study designs were popular, for a time, but were eclipsed as genome-wide association studies (GWAS) gained popularity [1]. GWAS have been effective for identifying population associations with common variants genome-wide, but have low power to study rare variants [2]. Family-based studies require smaller sample sizes than their case/control counterparts and enjoy increased power to detect effects of rare variants [2]. Additionally, family-based studies are able to identify next-generation sequencing (NGS) errors by utilizing familial relationships to identify unlikely calls [2]. Improvements in the cost and technology associated with NGS have facilitated a revival in family-based studies [1]. Family-based analyses coupled with NGS can uncover rare variants that are undetected by GWAS [2]. For example, analysis of whole exome sequence data was used to identify rare variants associated with non-syndromic oral clefts in large pedigrees ascertained to contain at least two affected relatives [3], to prioritize rare variants in large multi-generational pedigrees ascertained for multiple relatives diagnosed with bipolar disorder [4], and to identify rare variants segregating in families that contained at least two siblings with an autism spectrum disorder [5].

Unfortunately, family-based studies do not come without complication; for example, identifying a suitable number of pedigrees with desired criteria may be time consuming, sometimes requiring years to amass. In these circumstances, collecting new data to evaluate methodology or replicate findings is impractical. To address this challenge we have created an `R` package, entitled `SimRVPedigree`, which simulates pedigrees ascertained to contain a minimum number of disease-affected relatives. `SimRVPedigree` models the affected individuals in an ascertained pedigree as the result of (1) sporadic disease or (2) a single, rare, disease-variant segregating in the pedigree. At the individual level, `SimRVPedigree` models competing age-specific life events contingent on rare-variant status, disease status, and age through user supplied age-specific incidence rates of disease, and age-specific hazard rates for death. In a recursive manner, life events simulated at the individual level build and shape simulated pedigrees. Upon specification of user-defined study characteristics, `SimRVPedigree` will simulate pedigrees ascertained to contain multiple affected relatives according the specified criteria. To our knowledge, this is the only program to incorporate a competing risk model and account for the ascertainment process.

## Methods

Given a sample of pedigrees we allow for the possibility that different families may segregate different rare variants, but assume that within a family genetic cases are due to a shared rare variant that increases disease susceptibility. We allow users to choose between two methods of rare variant introduction to the pedigree. One option is to assume that all ascertained pedigrees with genetic cases are segregating a variant that is rare enough to have been introduced by exactly one founder [6]. Alternatively, we allow users to simulate the starting founder’s rare variant status with probability equal to the carrier probability of all causal variants considered as a group. When this option is selected some ascertained pedigrees may not segregate a causal variant. In either scenario, we assume that a causal variant is introduced by at most one founder and, when it is introduced, it is transmitted from parent to offspring according to Mendel’s laws.

Starting at birth and ending with death, we simulate life events for the starting founder, censoring any events that occur after the last year of the study. We repeat this process, recursively, for all descendants of the founder allowing life events at the individual level to shape successive generations of the pedigree. To accomplish this, we condition on an individual’s age, rare-variant status and disease status, and simulate waiting times to three competing life events: reproduction (i.e. producing offspring), disease onset, and death. We select the event with the shortest waiting time, update the individual’s age by this waiting time, record the event type, and repeat this process from the new age until the individual dies or the end of the study is reached.

### Simulating Life Events

To simulate life events `SimRVPedigree` users are required to specify: `hazardDF`, a data frame of age-specific hazard rates, where column one represents the age-specific hazard rates for the disease in the general population, column two represents the age-specific hazard rates for death in the **unaffected** population, and column three represents the age-specific hazard rates for death in the **affected** population, and `partition`, a discrete partition of ages over which to apply `hazardDF`.

Specifically, `partition` is a vector of ages, starting at age 0, such that `hazardDF[k,]` are the age-specific hazard rates for an individual whose age is contained in [`partition[k], partition[k+1]`). At the user’s discretion, if the disease of interest is rare, the age-specific hazard rates for death in the **unaffected** population may be approximated by age-specific hazard rates for death in the general population. In the following subsections, we detail the procedures to simulate waiting times to onset, death, and reproductive events.

#### Disease onset

We model disease onset using a non-homogeneous Poisson process (e.g. [7]), conditioned on an individual’s current age, *t*′, rare-variant status, *x*, and disease status, *δ*. In this context, *x* = 1 if the individual is a carrier of the rare variant, and 0 otherwise; and *δ* = 1 if the individual has developed disease by age *t*′, and 0 otherwise. Define *κ* to be the relative-risk of disease for individuals who have inherited the causal variant and *λ_o_*(*t*) to be the baseline age-specific hazard rate of disease for an individual aged t years. That is, *λ_o_*(*t*) is the age-specific hazard rate for individuals who do not carry a causal variant, i.e. sporadic cases. Let *λ_onset_*(*t*|*x*) denote the age-specific hazard rate of disease for an individual aged *t* years conditioned on rare-variant status such that

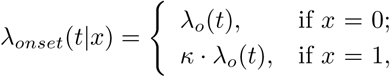

for *κ* ≥ 1.

If *p_c_* is the carrier probability of all causal variants considered as a group, then we can express the population age-specific hazard rate of disease, *λ_onset_*(*t*), as

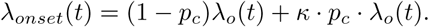

Users are expected to provide *λ_onset_*(*t*); given *p_c_* and *κ* we infer *λ_o_*(*t*) as *λ_o_*(*t*) = 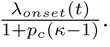 We note that this method for calculating *λ_o_*(*t*) has implications on the comparability of non-genetic individuals from studies simulated under very different *κ* values. For example, when *p_c_* is constant, we see that for *κ*_1_ ≪ *κ*_2_, the age-specific hazard rate for non-carrier individuals under genetic relative-risk *κ*_1_ will be much greater than that of non-carrier individuals under genetic relative-risk *κ*_2_. As *p_c_* increases this effect is visible more quickly for differing *κ* values.

We note that not all individuals develop the disease; however, those who do are only permitted develop the disease once in our model. Individuals who have developed disease (i.e. *δ* = 1) do not develop disease again, but can reproduce or die. When *δ* = 0, we use intensity function *λ_onset_*(*t*|*x*) conditioned on rare-variant status, *x*, to simulate the waiting time to disease onset given current age, *t*′. To clarify, if we denote the waiting time to disease onset by *W_onset_*, and condition on the current age, *t*′, the cumulative distribution function of *W_onset_* is given by

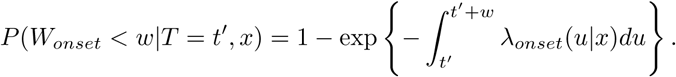

#### Death

We model death using a non-homogeneous Poisson process, conditioned on an individual’s current age, *t*′, and disease status, *δ*. Define *δ* as in the previous discussion, and let *λ_u_*(*t*) and *λ_a_*(*t*) denote the age-specific hazard rates of death, for individuals aged *t* years, in the **unaffected** population and the **affected** population, respectively. We use intensity function *λ_death_*(*t*|*δ*) conditioned on disease status *δ* to simulate the waiting time to death given the current age, *t*′. In this context, *λ_death_*(*t*|*δ*) represents the age-specific hazard rate of death for an individual aged *t* years conditioned on their disease status, which we model as

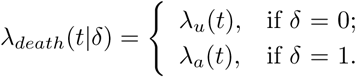

We do not model disease remission; after an individual has developed disease we use the age-specific hazard rates for death in the *affected* population to model their waiting time to death.

#### Reproduction

To accommodate extra-Poisson variability in the number of human offspring, we use a negative-binomial model with number of trials *n* ≈ 2 and success probability *p* ≈ 4/7, as proposed by [8]. We adopt this negative-binomial model of offspring number in `SimRVPedigree`. We employ an equivalent Poisson-Gamma mixture model [9] to obtain the negative-binomial offspring number and to simulate the waiting time to reproduction.

Let *w_t′_* denote the waiting time to reproduction given an individual’s current age *t*′, and assume that simulated subjects are able to reproduce from age *a_1_* to age *a*_2_. To mimic observed data on first-born live births (see `SimRVPedigree` Supplement, section 6), we simulate *a*_1_ and *a*_2_ as follows: sample a_1_ uniformly from ages 16 to 27, and *a*_2_ – *a*_1_ uniformly from 10 to 18 years. At birth we simulate an individual’s lifetime birthrate by taking a random draw, *γ*, from a gamma distribution with shape 2 and scale 4/3. Individuals who draw large *γ* will have high birth rates and many children, whereas individuals who draw small *γ* will have low birth rates and few or no children.

For some diseases, users may want to reduce the birth rate after disease onset; we allow users to achieve this through an additional parameter *f*, assumed to be between 0 and 1, which is used to rescale the birth rate after disease onset. By default, *f* = 1 so that the birth rate remains unchanged after disease onset. Given an individual’s birth rate, current age, and disease status, *δ*, we obtain their waiting time to reproduction as follows:

1. Simulate the unconditional waiting time to reproduction by drawing *w* from an exponential distribution with rate 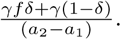
2. Condition on the current age, *t*′, to obtain the conditional waiting time to reproduction:

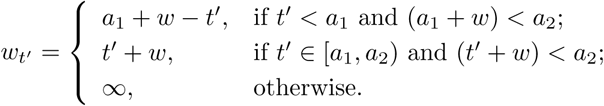

### Pedigree Simulation

To simulate all life events for a subject, starting at birth we generate waiting times to disease onset, death, and reproduction, as outlined previously and choose the event with the shortest waiting time to be the next life event. Next, we add the waiting time associated with the earliest event to the current age and either record the year of disease onset or death, or add a new offspring to the pedigree. We repeat this process from the updated age, recursively, until the individual dies or the study stop year is reached. This algorithm details the full life event procedure at the individual level. Complete details are available in an additional file [see `SimRVPedigree` Supplement, section 1].

To simulate a full pedigree, we recursively apply the algorithm described above, as follows:

- Step 1: Simulate life events for the first founder given rare-variant status.
- Step 2: Simulate life events for any new offspring given rare-variant status as outlined above.
- Step 3: Repeat step 2 until life events have been simulated for all offspring.

### Ascertainment Features

The primary function of `SimRVPedigree, sim_RVped()`, simulates pedigrees ascertained for multiple disease-affected relatives. We allow users to specify family-based study features through the following arguments of `sim_RVped()`:

`num_affected`: the minimum number of disease-affected relatives required for ascertainment of the pedigree.

`ascertain_span`: the start and stop year for pedigree ascertainment.

`stop_year`: the last year of follow-up for the pedigree.

`recall_probs`: the proband’s recall probabilities for relatives of varying degree.

In this context, the proband is the affected family member first in contact with the study, presumably at the time of disease onset.

The ascertainment span represents the time span, in years, over which the family could be ascertained through the proband. For example, suppose that a particular study ascertained families, containing at least two affected members, from 2000 to 2010. In this scenario, the user would set `ascertain_span = c(2000, 2010)` and `num_affected = 2`. The `sim_RVped()` function would then simulate families such that the proband developed disease between 2000 and 2010 and was at least the second family member to develop disease by then.

The study stop year represents the last year data are collected for ascertained families. Consider the previous study, and suppose that data were collected until 2016. To achieve this in simulation, users would simply specify `stop_year = 2016`, which would result in `sim_RVped()` simulating life events for ascertained families until the year 2016.

Often researchers involved in family-based studies are confronted by incomplete ascertainment of a proband’s relatives, which could occur if the proband cannot provide a complete family history, or if he or she does not support contact of specific relatives. `SimRVPedigree` allows users to mimic this scenario, in simulation, by trimming relatives from a pedigree based on the proband’s probability of recalling them. To specify a proband’s recall probabilities for his or her relatives, i.e. `recall_probs`, the user provides a list of length *q*, such as *p* = (*p*_1_, *p*_2_, …, *p_q_*). In this context, *p_i_* is used to denote the proband’s recall probability for a relative of degree *i* when *i* = 1, 2, …, *q* – 1, or the proband’s recall probability for a relative of degree *q* or greater when *i* = *q*. To simulate fully ascertained families, we set `recall_probs = c(1)`, which corresponds to *p* = 1. Alternatively, if unspecified, `recall_probs` is set to four times the kinship coefficient, e.g. [10]. This default value retains the proband’s first-degree relatives (i.e. parents, siblings, and offspring) with probability 1, second-degree relatives (i.e. grandparents, grandchildren, aunts, uncles, nieces, and nephews) with probability 0.5, third-degree relatives with probability 0.25, etc.

In the event that a trimmed relative is required to fully specify the relationships among recalled family members, we include the trimmed relative, mark them as unavailable, and remove (i.e. mark as missing) any of their relevant information. That is, disease status, relative-risk of disease, and event years are all missing for any relatives not recalled by the proband. Since disease-affected relatives may be trimmed from a pedigree, trimmed pedigrees may contain fewer than `num_affected` disease-affected relatives. When this occurs, `sim_RVped()` will discard the pedigree and simulate another until all conditions specified by the user are met.

## Results

### Settings

In the following applications, we use `SimRVPedigree` in conjunction with `R` [11] to investigate the effect of the relative-risk of disease in genetic cases, *κ*, on ascertained pedigrees. We first investigate the effect of *κ* on the number of affected relatives per family, and on the degree of familial clustering among affected relatives. Next, we investigate how ages of onset from more recent generations tend to be younger than those from older generations in the ascertained pedigrees [12], a phenomenon which we refer to as apparent anticipation. Lastly, we demonstrate how `SimRVPedigree` may be used to estimate the proportion of families that segregate the causal variant in a sample of ascertained pedigrees.

To study pedigrees ascertained to contain multiple relatives affected by a lymphoid cancer, we simulated study samples according to the following criteria.

1. Each study sample contained a total of one thousand pedigrees, ascertained from the year 2000 to the year 2015.
2. Each pedigree contained at least two relatives affected by lymphoid cancer.
3. The birth year of the founder who introduced the rare variant to the pedigree was distributed uniformly from 1900 to 1980.
4. For each *κ* considered, the carrier probability, *p_c_*, for all causal variants with genetic-relative risk *κ* was assumed to be 0.002.
5. Sporadic cases, i.e. affected individuals who did not inherit the rare variant, develop lymphoid cancer according to the baseline, age-specific hazard rate of lymphoid cancer. The population, age-specific hazard rate of lymphoid cancer were estimated through the Surveillance, Epidemiology, and End Results (SEER) Program [13, 14], and are displayed in Figure 1.
6. Genetic cases, i.e. affected individuals who did inherit the rare variant, develop lymphoid cancer at *κ* times the baseline, age-specific hazard rate of lymphoid cancer. We considered *κ* ∈ (1,10, 20) and simulated one thousand pedigrees for each *κ* considered.
7. Since lymphoid cancer accounts for a relatively small proportion of all deaths, the age-specific hazard rate for death in the unaffected population was approximated by that of the general population. Individuals who do not develop lymphoid cancer die according to the age-specific hazard rate of death in the general population [15], while individuals who have developed lymphoid cancer die according to the age-specific hazard rate of death in the affected population [13, 16, 17]. Figure 1 displays the age-specific hazard rates of death for these two groups.
8. The proband’s probabilities for recalling relatives were set to `recall_probs = (1, 1, 1, 0.5, 0.125)`, so that all first, second, and third-degree relatives of the proband were recalled with probability 1, all fourth-degree relatives of the proband were recalled with probability 0.5, and all other relatives of the proband were recalled with probability 0.125.
9. The stop year of the study was set to 2017.

**Figure 1.**
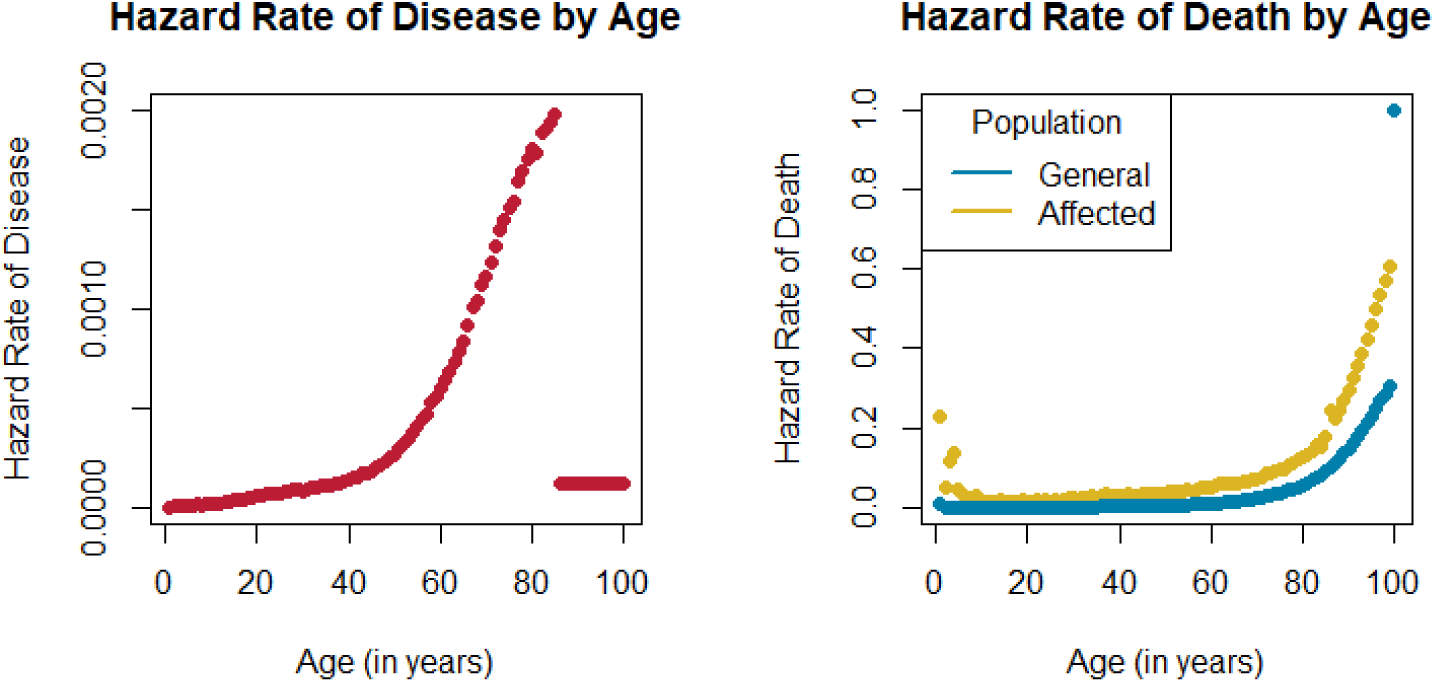
Hazard Rates. (Left) Baseline, age-specific hazard rates of lymphoid cancer estimated by SEER [13, 14]. SEER provides age-specific incidence and morality data, in yearly increments, up to age 84 years, and then aggregates data for ages of 85 years or greater. We considered the SEER reported incidence rate for individuals of age 85 or greater to be the constant hazard rate of disease for individuals between the ages of 85 to 100. (Right) Age-specific hazard rates of death for the general population [15] and for the disease-affected population [13, 16, 17]. To promote continuity in the age-specific hazard rate of death for the affected population, we assume that it is twice that of the unaffected population after age 84 years. After age 84 years, the SEER data do not allow for the age-specific hazard rates of death in the affected population to be estimated in yearly increments.

### Example

We demonstrate how to simulate a single pedigree according to the settings described previously.

After installing `SimRVPedigree`, we load the package in `R` using the `library` function.

`R> library(SimRVPedigree)`

Suppose that we can obtain age-specific hazard rates in yearly increments starting at age 0 and ending with age 100. In this case, we define the partition of ages over which to apply the age-specific hazards rates using the `seq` function.

`R> age_part <-seq(0, 100, by = 1)`

Next, assume that `LC_Hazards` is a data frame whose columns provide age-specific hazard rates, in yearly increments, from age 0 to age 100, as indicated below.

`LC_Hazards[, 1]` Age-specific hazard rates of lymphoid cancer in the general population.

`LC_Hazards[, 2]` Age-specific hazard rates of death for individuals in the general population.

`LC_Hazards[, 3]` Age-specific hazard rates of death for individuals who have lymphoid cancer.

We create a new object of class hazard from the partition of ages, `age_part`, and the data frame of hazard rates, `LC_Hazards`, by executing the following command.

~~~
R> haz_mat <-hazard(partition = age_part,
                    hazardDF = LC_Hazards)
~~~

To simulate a single pedigree with family identification number 1 and a genetic relative-risk of 10, assuming that the eldest founder introduces the variant, and according to the settings described previously in *Results: Settings* we use the following command.

~~~
R> ex_ped <-sim_RVped(hazard_xates = haz_mat,
                      GRR = 10, FamID = 1,
                      num_affected = 2,
                      RVfounder = TRUE,
                      ascertain_span = c(2000, 2015),
                      founder_byears = c(1900, 1980),
                      recall_probs = (1, 1, 1, 0.5, 0.125),
                      stop_year = 2017)
~~~

To view a description of the contents of `ex_ped` we use the `summary` command.

~~~
R> summary(ex_ped)
                 length   Class   Mode
 full_ped      15    ped   list
 ascertained_ped     15     ped   list
~~~

Upon executing the command above, we see that `ex_ped` is a list containing two objects of class `ped`. The first is named `full_ped` and represents the original pedigree, prior to proband selection and trimming. The second is named `ascertained_ped` and represents the ascertained pedigree; this data frame includes an additional variable to identify the proband. In this application, we are interested in families that were ascertained for study; hence, we focus attention on `ascertained_ped`.

To simplify the following examples, we store the ascertained pedigree as `study_ped`.

`R> study_ped <-ex_ped$ascertained_ped`

To plot the ascertained pedigree we simply supply the pedigree to the `plot` function.

`R> plot(study_ped)`

The plotted pedigree is displayed in Figure 2.

**Figure 2.**
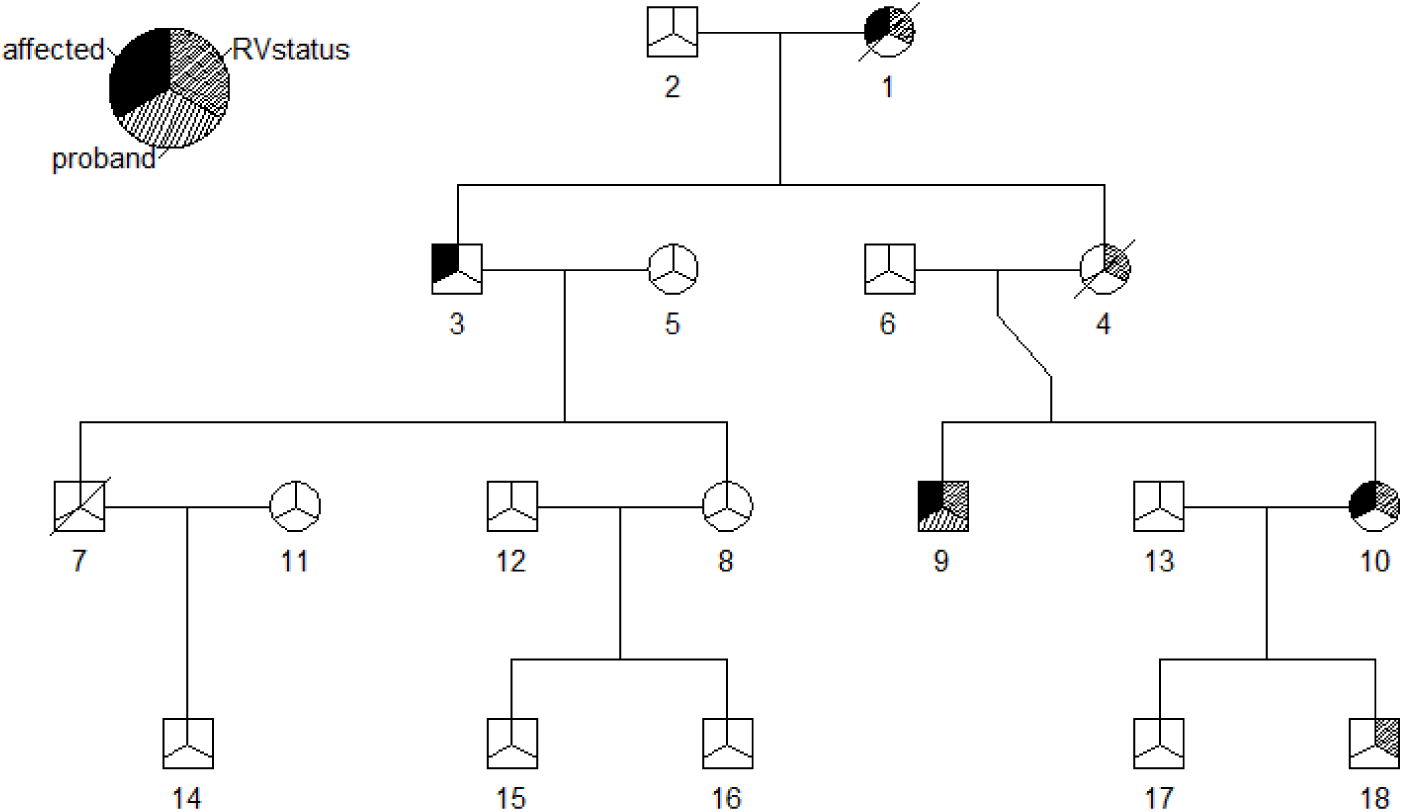
Simulated Pedigree. In this pedigree squares are used to symbolize males and circles are used to symbolize females. Mates are connected by a horizontal line, and their offspring branch out below. Individuals who have died have a slash through their symbol. As indicated by the legend, if the upper left third of an individual’s symbol is shaded black, then that individual is disease-affected. If the upper right third of an individual’s symbol is shaded, then that individual is a carrier of the causal variant. If the bottom third of an individual’s symbol is shaded, then that individual is the proband.

To obtain summary information for `study_ped` we supply it to `summary`.

~~~
R> summary(study_ped)
$family_info
 FamID  total_rels   num_affected   ave_onset_age   ave_IBD   asc_year   seg_RV
     1          18          4              52.5       0.333     2002      TRUE
$affected_info
 FamID   ID   birthYr   onsetYr   deathYr   RR   proband   RVstatus
     1    1   1911      1965        1968    10     FALSE    1
     1    3   1933      2014         NA      1     FALSE    0
     1    9   1966      2002         NA     10      TRUE    1
     1   10   1972      2011         NA     10      FALSE   1
~~~

As displayed above, when the argument of `summary` is an object of class `ped, summary` returns two data frames named `family_info` and `affected_info`. The `family_info` data frame catalogues the information for the entire family. For each family supplied it provides (from left to right): family identification number, the total number of relatives in the pedigree, the total number of disease-affected relatives in the pedigree, the average onset age of the disease-affected relatives, the average of the pairwise probabilities of identity by descent (IBD) among the disease-affected relatives in the pedigree, the ascertainment year of the pedigree, and a logical variable indicating whether or not the pedigree segregates a casual variant. The `affected_info` data frame catalogues information for the disease-affected relatives. For each disease-affected relative it details (from left to right): family identification number, individual identification number, year of birth, year of disease-onset, year of death, relative risk of disease, proband status, and rare variant status.

### Applications

#### Number of Disease-Affected Relatives

To illustrate how the number of disease-affected relatives in each pedigree varies with *κ*, we refer to the data described in *Results: Settings*. This data contains simulated study samples, containing 1000 pedigrees, for *κ* = 1, *κ* = 10, and *κ* = 20.

Figure 3 summarizes the distribution of the number of disease-affected relatives per pedigree for these three groups. From the figure we see that for *κ* = 1 this distribution is more highly concentrated at two affected members than for the other two groups considered. Not surprisingly, as *κ* increases we see relatively fewer families containing only two affected members, and more families containing three or greater affected members.

**Figure 3.**
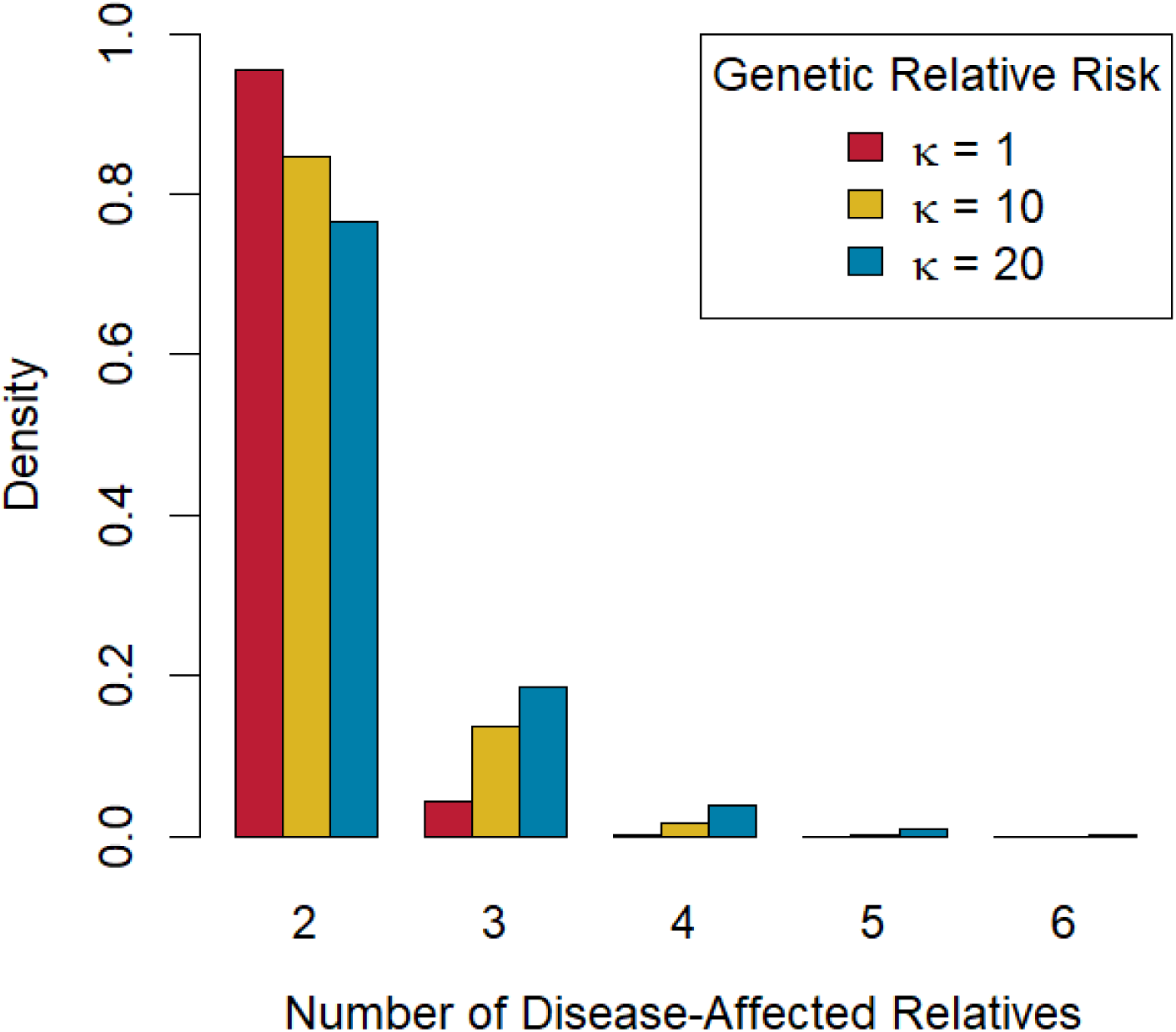
Bar charts of Number of Disease-Affected Relatives per Pedigree. Barcharts of number of disease-affected relative per pedigree grouped by genetic relative-risk of disease, *κ*.

#### Familial Clustering

To investigate the relationship between familial clustering among affected relatives and *κ*, we restrict attention to pedigrees that contained two or three affected relatives. We did not consider pedigrees with four or more disease-affected relatives because these pedigrees are rarely observed when *κ* = 1. This resulted in a total of 999 simulated pedigrees in the *κ* = 1 group, 982 simulated pedigrees in the *κ* = 10 group, and 950 simulated pedigrees in the *κ* = 20 group. To assess the level of familial clustering among affected relatives, we computed the average of the pairwise IBD probabilities among affected members in a pedigree, which we will denote by 𝓐_*IBD*_. 𝓐_*IBD*_ is proportional to the genealogical index of familiality statistic [18], which has been used to summarize familial clustering of aggressive prostate cancer in the Utah population. In general, the IBD probability between two relatives decreases as they become more distantly related. For example, for an affected parent-child pair, or two affected siblings 𝓐_*IBD*_ = 0.5; whereas for an affected avuncular pair, or an affected grandparent-grandchild pair 𝓐_*IBD*_ = 0.25.

Figure 4 shows the conditional distribution of 𝓐_*IBD*_ given the total number of affected relatives in a pedigree and *κ*. Tabulated results for Figure 4 are available in an additional file [see `SimRVPedigree` Supplement, section 2]. The left panel of Figure 4 summarizes the conditional distribution of 𝓐_*IBD*_ for families with two affected members. The conditional distribution of 𝓐_*IBD*_ shifts probability mass toward 0.5 as *κ* increases and suggests that disease-affected individuals tend to be more closely related in families with larger values of *κ*. The right panel of Figure 4 summarizes the conditional distribution of 𝓐_*IBD*_ among families with three affected members, and shows the same trend as the left panel, of 𝓐_*IBD*_ values shifted towards 0.5 for larger values of *κ*.

**Figure 4.**
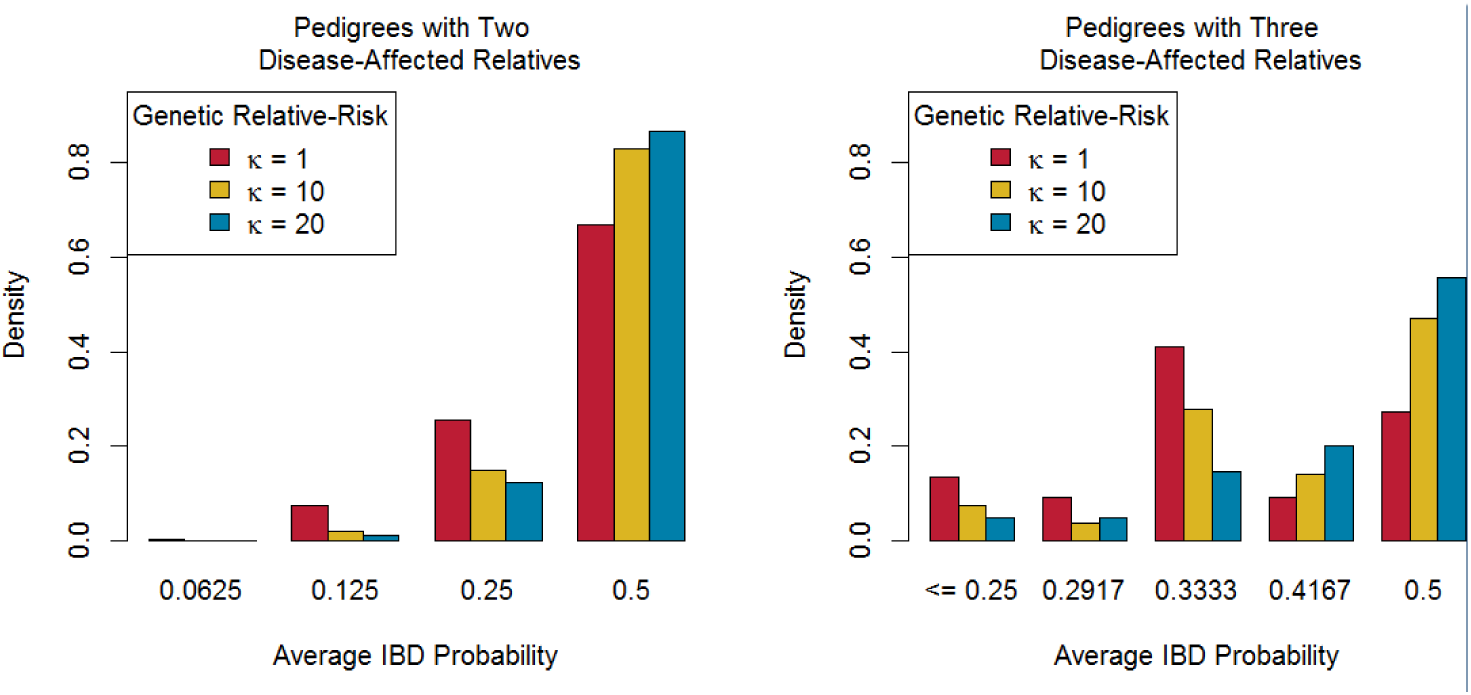
Bar charts of 𝓐_*IBD*_ Distributions. Barcharts of 𝓐_*IBD*_ distributions for pedigrees with two (left) or three (right) disease-affected relatives, grouped by genetic relative-risk of disease.

#### Anticipation

Anticipation is a decreasing trend in the age of disease onset, and possibly an increasing trend in severity, in successive generations of a family [19]. Some genetic diseases with unstable repeat expansions show anticipation, and include: Huntington’s Disease, fragile X syndrome, and myotonic dystrophy [20].

However, studies of genetic anticipation based solely on the ages of onset of affected members have the potential for ascertainment bias [21]. Possible sources of ascertainment bias include: early detection in offspring due to parental diagnosis or improved diagnostic techniques and right-censoring of family members who have developed the disease by the end of the study, especially in studies of large multi-generational pedigrees that have been ascertained to contain multiple affected members. [12, 21].

Referring to the data described in section *Results: Settings*, we illustrate how apparent anticipation can arise as an artefact of studies ascertaining families with multiple disease-affected relatives. Within each of the families considered, generation number was assigned among affected relatives so that generation number one represents the most recent common ancestor with whom all affected members could share a variant identical by descent. In this assignment scheme, we allow an affected individual to be his or her own most recent common ancestor. To demonstrate this convention, consider a family with two affected relatives: if the affected members are a parent-child pair, then the parent would be assigned generation number one, and the child assigned generation number two. However, if the affected members are a sibling pair, each sibling would be assigned generation number two, since a parent is the closest relative from whom the affected siblings could have inherited a disease variant.

Figure 5 displays the ages of onset, by assigned generation, grouped by *κ*, the relative-risk of disease for genetic cases. We emphasize that `SimRVPedigree` does not include a mechanism to simulate anticipation. However, we note that even though anticipation is not present in the simulated data, within each genetic-relative-risk group considered, the box plots exhibit a decreasing trend in the ages of onset for successive generations. The false anticipation signal is likely due to many of the ascertained pedigrees being large, and multi-generational, and therefore prone to right-censoring of younger family members who will develop disease later in life, after the study stop year.

**Figure 5.**
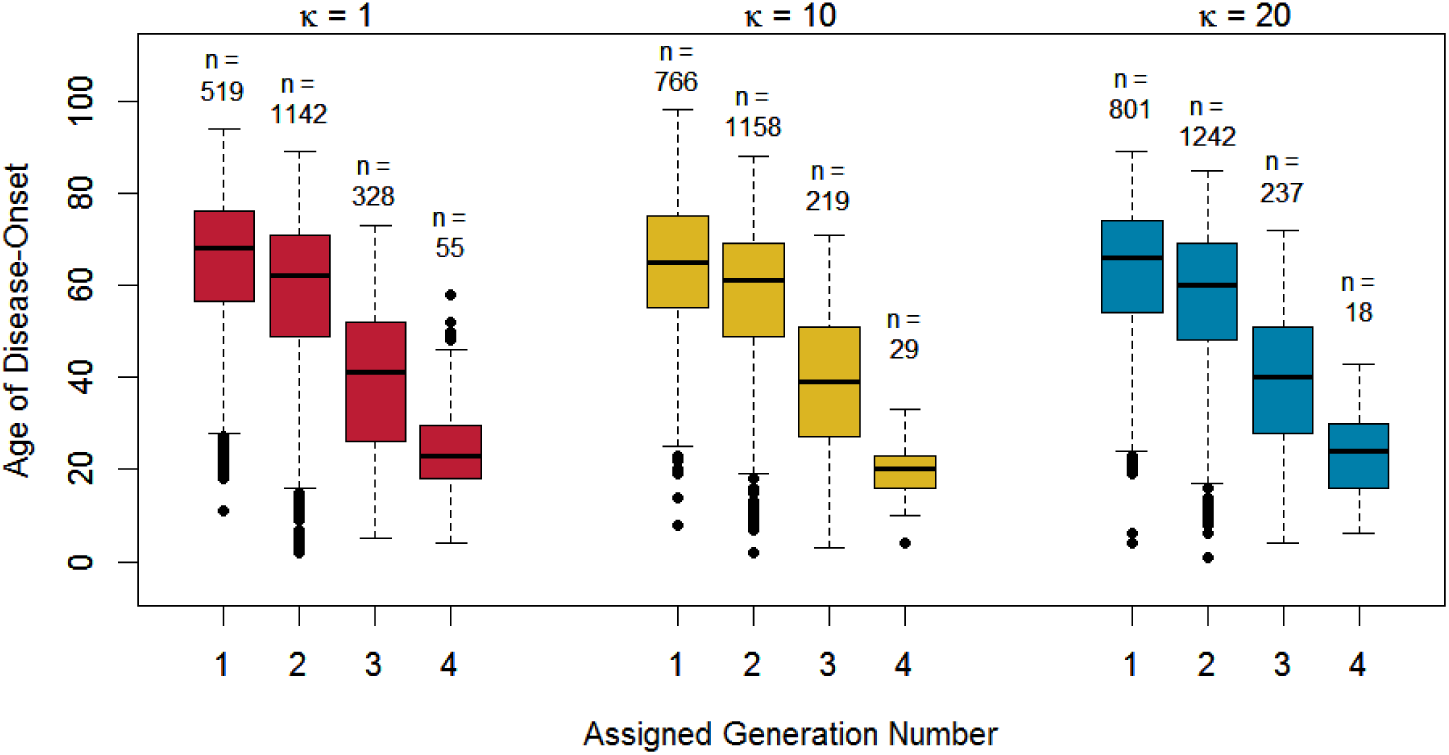
Box plots of Age of Disease Onset by Assigned Generation Number. Boxplots of age of onset by assigned generation number, as defined in text, grouped by genetic relative-risk of disease, *κ*. The numbers of observations, *n*, used to create each box plot are displayed above their respective plots.

If there is right-censoring of younger family members then this censoring should be apparent in their ages of death as well. Therefore it is useful to consider using the ages of death in unaffected relatives as a negative control to gain insight into ascertainment bias [19]. Box plots of the ages of death in unaffected relatives by generation for the relative-risk groups are similar to those in Figure 5 for the age of onset in disease-affected relatives. This similarity strongly suggests the presence of ascertainment bias. Further details of this investigation may be found in an additional file [see `SimRVPedigree` Supplement, section 2].

#### Proportion of Ascertained Pedigrees Segregating a Causal Variant

Familial lymphoid cancer, i.e. a family that contains multiple relatives affected by lymphoid cancer, is relatively rare; however, lymphoid cancer is not a rare disease as it affects roughly 1 in 25 [13, 14]. With such diseases, there is a greater risk of ascertaining pedigrees that contain multiple disease-affected relatives by chance alone. Since we do not expect these pedigrees to segregate a causal variant it is advantageous to choose ascertainment criteria that reduces the likelihood of sampling such pedigrees.

To determine what proportion of ascertained families we expect to segregate a causal variant we conducted a simulation study in which the rare variant status of the starting founder was allowed to vary so that fully sporadic pedigrees were given an opportunity for ascertainment.

The procedure to simulate a study containing both genetic and sporadic families may be described as follows.

Step 1: Allow the starting founder to introduce a causal variant with genetic relative-risk *κ* with probability 0.002.
Step 2: Simulate the rest of the pedigree, according to the settings described in *Results: Settings*, and add it to our sample of ascertained pedigrees if it meets the ascertainment criteria.
Step 3: Repeat steps one and two until the requisite number of pedigrees have been ascertained.

For this procedure we considered *κ* = 1 and all multiples of 5 between 5 and 100, i.e. *κ* ∈ *c*(1, 5,10,15,…, 95,100). For each *κ* considered we simulated family studies each containing one thousand ascertained pedigrees. Next, we determined what proportion of the ascertained pedigrees were segregating a causal variant that increased disease susceptibility. The results of this investigation are displayed in Figure 6. The leftmost panel in Figure 6 indicates that most of the ascertained pedigrees are not segregating a causal variant. For example, when the genetic relative-risk is 20, we see that less than 20% of the ascertained pedigrees with two or more disease-affected relatives are segregating a causal variant. Focusing attention on the ascertained pedigrees that contain three or more affected relatives (the middle panel of Figure 6) we see that these pedigrees tend to segregate a causal variant more often than the pedigrees that only contained two or more affected relatives. When we restrict our focus to the ascertained pedigrees that contain four or more affected relatives (the rightmost panel of Figure 6), we see more of these pedigrees tend to segregate a causal variant. These estimates tend to be more erratic because we don’t often observe fully sporadic families with four or more affected relatives. We did not observe fully sporadic pedigrees with five or more affected relatives in any of the original samples of one thousand pedigrees.

**Figure 6.**
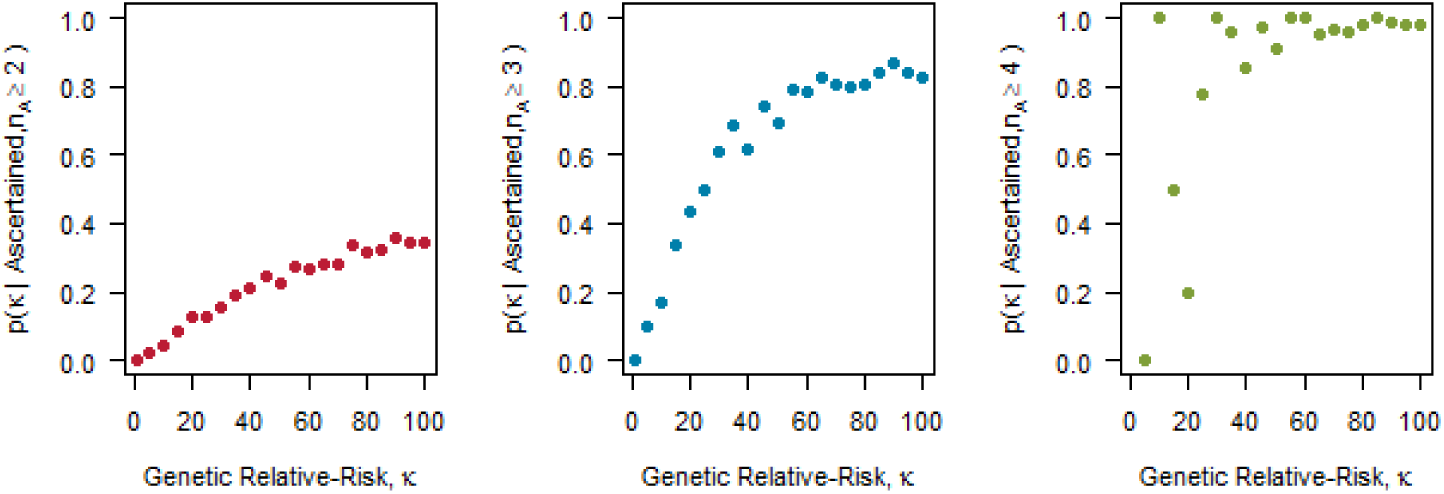
Genetic Contribution Estimate. Scatter plots of the probability that a randomly selected pedigree from a sample of ascertained pedigrees is segregating a genetic variant with relative-risk of disease *κ* against the relative-risk of disease *κ*. Here we consider the effect of restricting attention to the ascertained pedigrees with *n_A_* or more disease-affected relatives. In the leftmost panel, we consider all one thousand pedigrees ascertained with two or more disease-affected relatives; in the middle panel, we consider the subset with three or more disease-affected relatives, and in the right most panel the subset with four or more disease-affected relatives.

These results indicate that when a disease is not rare, and when the carrier probability of the causal variant is very low (i.e. *p_c_* = 0.002), focusing on families with at least three affected relatives is more effective for sampling pedigrees that segregate a causal variant. Focusing on pedigrees with at least four affected relatives provides even greater improvement.

#### Computation Time

We would like to note that simulation of ascertained pedigrees can be computationally expensive. Therefore, we urge users to take advantage of parallel processing, in R, or cluster computing when simulating a large number of ascertained pedigrees.

There are several factors that effect the amount of time required to simulate a pedigree. For example, the genetic relative-risk, the probability that a causal rare variant is segregating in the family, and the ascertainment span, to name a few. To illustrate the effect of the genetic relative-risk on timing we consider the family-study described in *Results: Settings.* The following table provides summary statistics for the average computation time, in seconds, required to simulate a single pedigree on a Windows OS with an i7-4790 @ 3.60 GHz, 12 GB of RAM, and a C220 SATA AHCI.

**Table 1.** Comparison of Computation Time for Various Genetic Relative-Risk Values. Tabulated average computation time and standard deviation of computation time, in seconds. These results were obtained over 50 repeated simulations of a single pedigree.

| Genetic Relative-Risk | Average Simulation Time (in seconds) | Standard Deviation of Simulation Time (in seconds) | Number of Trials |
| --- | --- | --- | --- |
| 1 | 30.752 | 32.539 | 50 |
| 10 | 1.461 | 1.339 | 50 |
| 20 | 0.650 | 0.564 | 50 |

When probability that a causal rare variant is segregating in the family is small, the simulation time will tend towards the time required to simulate an ascertained pedigree with a genetic relative-risk of 1. This is the case for all pedigrees simulated in *Results: Proportion of Ascertained Pedigrees Segregating a Causal Variant* since the probability that the eldest founder introduces the rare variant is 0.002.

## Discussion

We provide several applications for `SimRVPedigree` to illustrate the effect of the genetic relative-risk, *κ*, on features of the ascertained pedigrees. First, we investigate the relationship between *κ* and the number of affected individuals in each ascertained family. In this application, as *κ* increases we observe pedigrees that contain three or more affected relatives more frequently than pedigrees with only two affected relatives.

Second, we examine the relationship between *κ* and the average, pairwise IBD probability among affected relatives in a pedigree. We observe that pedigrees simulated with larger values of *κ* tend to contain affected relatives that are more closely-related than pedigrees simulated with smaller values of *κ*.

Third, we illustrate that the family-based study design can contribute to apparent anticipation signals. In part, this is due to large, multi-generational pedigrees, which are prone to right-censoring of younger family members likely to experience disease onset later in life. This type of right-censoring can confound true genetic anticipation. We observe that it is possible to reduce this bias by following family members available at the time of ascertainment for a sufficient length of time. However, the necessary time frame (roughly 100 years) is impractical for real studies [see `SimRVPedigree` Supplement, section 5].

Finally, we show how users can estimate the proportion of ascertained pedigrees that are segregating a variant that increases disease susceptibility. In this application we find that when the carrier probability of all causal variants considered as a group is 0.002, many of the pedigrees ascertained with two or more disease-affected relatives do not segregate a genetic variant. In this scenario, it may be advantageous for researchers to focus on pedigrees with three or more disease-affected relatives. We note that when the carrier probability increases results will vary [see `SimRVPedigree` Supplement, section 6]. `SimRVPedigree` is intended for simulating diseases that are influenced by rare variants (e.g. allele frequency ≤ 0.005); however, when the carrier probability is increased to reflect variants that are less rare (e.g. allele frequency ∈ [0.005, 0.01]), `SimRVPedigree` may underestimate the proportion of ascertained pedigrees that contain genetic cases.

We emphasize that ascertained families can differ substantially depending on the simulation settings chosen. For example, variations in the ascertainment span can affect the distribution of the number of affected relatives in each pedigree, when all other study settings remain constant.

## Conclusions

The `SimRVPedigree` package provides methods to simulate pedigrees that contain multiple disease-affected relatives by a family-based study. To simulate life events at the individual level, `SimRVPedigree` models disease onset, death, and reproduction as competing life events; thus, pedigrees are shaped by the events simulated at the individual level. `SimRVPedigree` allows for flexible modelling of disease onset through user-supplied age-specific hazard rates for disease onset and death, and also permits flexibility in family-based ascertainment.

Among their benefits, family-based studies of large pedigrees with multiple disease-affected relatives enjoy increased power to detect effects of rare variants [2]. However, to conduct a family-based study of a rare disease it may take years to collect enough data. For planning and inference, we present the `SimRVPedigree` package to readily simulate pedigrees ascertained for multiple relatives affected by a rare disease. To our knowledge, this is the first package to dynamically simulate pedigrees to account for competing life events.

## List of Abbreviations

GWAS: genome-wide association studies
IBD: identity by descent
NGS: next-generation sequencing

## Declarations

### Ethics approval and consent to participate

Not applicable.

### Consent for publication

Not applicable.

### Availability of data and materials

`SimRVPedigree` is a platform-independent package and is readily available for `R` ≥ 3.4.0 through the CRAN repository. `SimRVPedigree` requires `kinship2` ≥ 1.6.4 and `dplyr` ≥ 0.7.4.

The datasets generated and/or analysed during the current study are available in the SimRVPedigree—Supplementary-Data repository, https://github.com/cnieuwoudt/SimRVPedigree—Supplementary-Data.

## Footnotes

**Author’s contributions** CN developed the R package, drafted the manuscript and conducted the analyses. SJ and ABW provided the initial motivation for the project and genetics guidance. JG conceived of the R package and the overall statistical approach, and contributed to the development of the manuscript and the applications. All authors read and approved the final manuscript.

## Acknowledgements

Not applicable.

## Additional Files

### Additional file 1 — SimRVPedigree Supplement

This is a pdf file that provides detailed information about the simulation procedure, as well as additional information for the applications discussed in the main text.

